# Bbs5 functions as the docking compartment of Bbsome binding to Klc3 regulated by Bbs5-Ser^246^ PKC-phosphorylation during mouse spermatogenesis

**DOI:** 10.1101/2020.03.06.981308

**Authors:** Dahai Yu, Lixia He, Xin Zhou, Xiuxia Wang, Bingzhi Yu

## Abstract

A mitochondrial and a fibrous sheath form the midpiece of the mammalian sperm flagellum encircling most of the axoneme. It has been documented that Kinesin light chain 3 (KLC3) was involved although the formation procedure remains unclear. Yeast-two-hybrid dataset showed an interaction between Klc3 and Bardet-Biedl Syndrome 5 (BBS5) Protein, another molecular associated with cilia and flagella forming. In this study, we presumed that the most conserved IFT complex BBsome was involved in spermatogenesis via the interaction of one of its subunits, Bbs5 with Klc3. Firstly, the interaction between Klc3 and Bbs5 was confirmed with Co-IP. Secondly, we identified PKC phosphorylation sites *in vitro* by LC-MS/MS, Ser^19^ and Ser^246^ of Bbs5, examined the phosphorylation status of Bbs5 Ser^19^ and Ser^246^ in mouse testis. Co-IP was performed to find which PKC isoforms phosphorylate Bbs5. In addition, we tried to discuss the roles of Ser^19^ and Ser^246^ of Bbs5 in the Klc3-bbs5 interaction and in mouse spermatogenesis based on our early findings.

## Introduction

The eukaryotic flagellum, a highly specified organelle, is responsible for the propulsion of the male gamete in most animals. A mitochondrial and a fibrous sheath form **the midpiece of the sperm** flagellum encircling most of the axoneme. Kinesins are a family of proteins, essential for the proper function of many polar cells, including epithelial cells, neurons, and sperm. Spermatogenesis is closely associated with many different kinesins [1]. Kinesin light chain 3 (KLC3) is the only known kinesin light chain expressed in post-meiotic male gamete [2]. Data show that spermatids KLC3 binds to outer dense fibers and mitochondrial sheath of midpiece of sperm tail [2, 3], and low expression of KLC3 might be associated with oligozoospermia and asthenozoospermia [4]. But a lot of details remain unknown.

The BBSome, consisting of BBS1, BBS2, BBS4, BBS5, BBS7, BBS8, BBS9, and BBS18 (also known as BBIP1) is a highly conserved protein complex regulating IFT in cilia and flagella, and mutations of BBSome subunits cause Bardet-Biedl Syndrome (BBS) [5–14]. In our previous study, BBS5 assembles into the BBsome as a peripheral, non-framework subunit [10]. Novel BBs5 mutations have been eventually found but its actual function remains very limited understood. Recent studies indicated that, in retina, Bbs5/Bbs5L (a newly found splice variant) localize at axoneme via binding to arrestin1 in a phosphorylation-dependent manner, and BBs5/Bbs5L are phosphorylated by protein kinase C (PKC) [15, 16]. The Bbs5 structural properties in BBsome and the Bbs5-arrestin1 binding indicated that Bbs5 might be functioning as a docking compartment of BBsome, and the covalent modifications of phosphorylation might be a good entry point to investigate the BBS5 functions. Meanwhile, no data has been show about the PKC phosphorylation sites of Bbs5. As a pivotal role in various biological processes, PKC also takes part in spermatogenesis [17–19]. Bbs5 might be one target how PKC get involved in since BBS5 is the subunit of BBsome, an indispensable complex during ciliogenesis and spermatogenesis [20, 21]. We predicted Bbs5 as a potential PKC substrate in mice with the DISPHOS software [22], and two potential PKC phosphorylation sites, including Ser^19^ and Ser^246^, were predicted.

A systematic proteome-scale mappping of ~14,000 binary protein-protein interactions has inspired us in further functional investigation on both Klc3 and Bbs5. Yeast-two-hybrid dataset indicated an interaction between Klc3 and Bbs5 [23] although a confirmation is still needed. Are this interaction co-related with spermatogenesis? If yes, what is regulating this interaction? Would the 2 “predicted” phosphorylation sites take some kind of roles in it? This study was trying to address these presumptions. Initially, we began with the Co-IP to confirm the Klc3-Bbs5 interaction as a supplement of yeast-two-hybrid result [23]. Then, for the first time, by LC-MS/MS analysis following *in vitro* phosphorylation, we identified the PKC phosphorylation sites including Ser^19^ and Ser^246^ of Bbs5, consistent with the DISPHOS prediction. Additionally We found the various phosphorylation statuses of Ser^19^ and Ser^246^ of Bbs5 in mouse brain and testis with customized phospho/nonphospho-Bbs5(Ser19) and phospho/nonphospho-Bbs5(Ser246) antibodies, showing that Ser^19^ is phosphorylated in both brain and testis, while Ser^246^ is only phosphorylated in brain, not in testis. Then we proved that Bbs5 is the direct downstream substrate of several dominant PKC isoforms in mouse testis. At this point, we seemingly have some confidence that Ser^19^ and Ser^246^ are certain targets of PKC, and potentially regulate of Klc3-Bbs5 interaction. The phosphorylation/dephosphorylation of Bbs5 Ser^19^ and Ser^246^ were mimicked. Consistent with the expression data, different combinations of mimicking phospho/dephospho mutants show various effects on Klc3-Bbs5 interaction, indicating that Ser^246^ phosphorylation/dephosphorylation may be the testis-specific switch regulating Klc3-Bbs5 interaction. Then as a supplement, more factors were introduced in: several constructs with known pathogenic mutations were built up; PKC was activated/deactivated by stimulator/inhibitor in media with or without serum, to observe the effects on Klc3-Bbs5 interaction.

### Ser^19^ and Ser^246^ Sites of Bbs5 Protein Are Phosphorylated by PKCs in Vitro

The results of LC-MS/MS analysis of phosphorylated Bbs5 were shown in Fig. 2, *A*. Two PKC phosphorylation sites, Ser^19^ and Ser^246^, were identified.

### Bbs5 is the direct downstream substrate of various PKC isoforms in mouse testis, Bbs5 Ser^19^ and Ser^246^ are both phosphorylated in brain, while only Se19^6^ is phosphorylated in testis

To identify whether Bbs5-Ser^19^ was phosphorylated *in vivo*, we collected mouse brain and testis tissue then measured the phosphorylation status of Bbs5-Ser^19^ and Ser^246^ by the phosphospecific antibodies, and the nonphosphorylation status by the nonphosphospecific antibodies. Fig.3 *A* showed that phosphorylated Bbs5-Ser^19^ band was observed in both brain and testis, whereas stronger phosphorylation of Bbs5-Ser^19^ was observed. Interestingly, a phosphorylated Bbs5-Ser^246^ band was detected in brain but absent in testis. Taken together, these results demonstrated that both Bbs5-Ser^19^ and Bbs5-Ser^246^ are phosphorylated in brain while only Bbs5-Ser^19^ in testis, indicating a tissue specificity of Bbs5 phophorylation. Also, we are interested in which PKC isoforms interact with Bbs5 in testis and the interactions were confirmed between 5 dominant PKC isoforms in testis, PKC α, β, γ, δ, ζ and Bbs5.

### Simulated phosphorylation/nonphosphorylation on single Ser site or both Ser sites regulate the Bbs5-Klc3 interaction

6 site-directed mutantgenesis Bbs5 constructs were built up, with Simulated phosphorylation/nonphosphorylation mutations on Ser^19^, Ser^246^ or both Ser^19^/Ser^246^. Simulated phosphorylation/nonphosphorylation on Ser^19^ have no effect on Bbs5-Klc3 interaction (Fig. 4 A.), indicating this interaction an intrinsic property of Bbs5 binding to Klc3; Interestingly, simulated phosphorylation on Ser^246^ completely block Bbs5-Klc3 interaction regardless of Ser^19^ (Fig. 4 B, C.), consistent with the Endogenous Bbs5 expression data.

### Some confirmed pathogenic Bbs5 mutations do not interrupt the Klc3-Bbs5 interaction

Additionally, We tried to observe whether several known BBs5 mutations could affect this interaction [23–25]. Only the pathogenic missense mutants were picked up for making the corresponding constructs. Co-IPs were performed and the data shows that none of mutations could 100% block the Klc3-Bbs5 interaction, while only G72S show a weakened pulled-down. More attention would be paid on in future study.

**Table 1.**
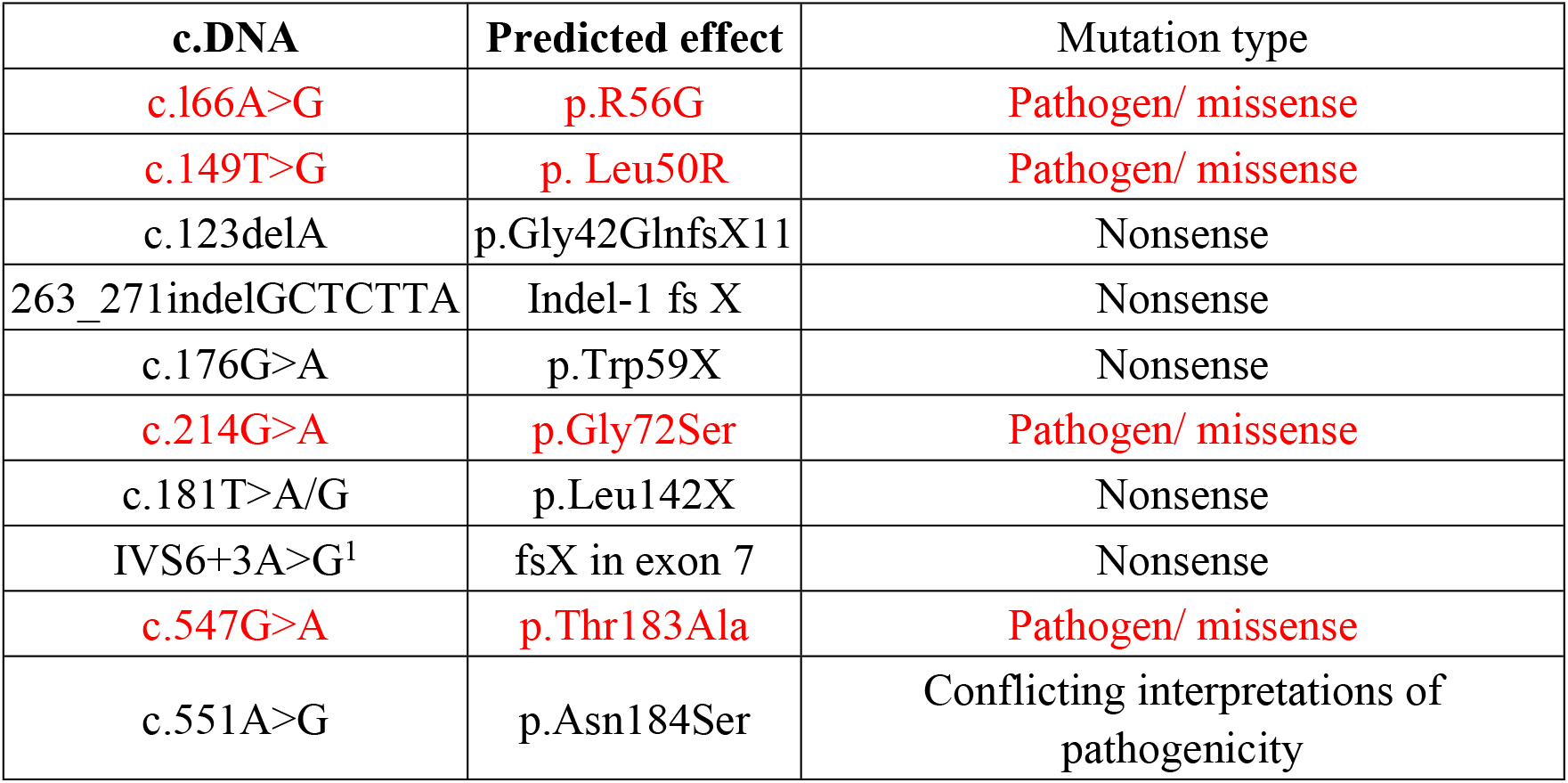
Mutations Reported in *BBS5*.

| c.DNA | Predicted effect | Mutation type |
| --- | --- | --- |
| c.166A>G | p.R56G | Pathogen/ missense |
| c.149T>G | p. Leu50R | Pathogen/ missense |
| c.123delA | p.Gly42GlnfsX11 | Nonsense |
| 263_271indelGCTCTTA | Indel-1 fs X | Nonsense |
| c.176G>A | p.Trp59X | Nonsense |
| c.214G>A | p.Gly72Ser | Pathogen/ missense |
| c.181T>A/G | p.Leu142X | Nonsense |
| IVS6+3A>G<sup>1</sup> | fsX in exon 7 | Nonsense |
| c.547G>A | p.Thr183Ala | Pathogen/ missense |
| c.551A>G | p.Asn184Ser | Conflicting interpretations of pathogenicity |

### PKC stimulation/inhibition interrupt the Klc3-Bbs5 interaction

All previous results come to that the Klc3-Bbs5 binding in testis is more acting like the intrinsic assembly under the various phosphorylation status combination of Ser19 and Ser246. We also want to understand what if the PKC was just stimulated or inhibited. The Klc3-Bbs5 binding was observed again under several different treatments. In the medium with/without serum, the PKC stimulator TPA or inhibitor H7 was applied to up-regulate or down-regulate the PKC activity. Apparently, serum starvation itself does not block the binding. Surprisingly, both the stimulation and the inhibition of PKC would almost 100% block the interaction.

## DISCUSSION

Eukaryotic flagella and cilia share a basic 9+2 microtubule-organization of axonemes, and could generate bending-motion [28]. The eukaryotic flagellum, generally considered as a highly specified cilium, is responsible for the propulsion of the male gamete in most animals. Compatible with this function, a mitochondrial and a fibrous sheath has been evolved to form **the midpiece of the sperm** flagellum, encircling most of the axonemes. Kinesins are a family of proteins, essential for the proper function of many polar cells, including epithelial cells, neurons, and sperms. Spermatogenesis is closely associated with many different kinesins [1]. Kinesin light chain 3 (KLC3), a member of the kinesin-1 family, is the only known kinesin light chain expressed in postmeiotic male gamete [2]. It contains a conserved heptad repeat (HR) binding to kinesin heavy chain (KHC) [29, 30] and tandem tetratrico-peptide repeats (TPR) mediating protein interactions in [31]. KLC3 are highly expressed in mouse round and elongating spermatids. KLC3 accumulates in the sperm tail midpiece consisting of a mitochondrial sheath [32]. KLC3 binds to mitochondria during rat spermatogenesis and was believed functioning either as a vehicle or a fixation [33]. Low expression of KLC3 might be associat6hged with oligozoospermia and asthenozoospermia [4]. KLC3 is proposed to play multiple roles in sperm tail development. But a lot of details remain unknown.

BBS5 is one of the known 20 pathogenic genes causing Bardet-Biedl Syndrome (BBS) [5–14]. The BBSome, consisting of BBS1, BBS2, BBS4, BBS5, BBS7, BBS8, BBS9, and BBS18 (also known as BBIP1) is a highly conserved protein complex regulating IFT in cilia and flagella. In my sequential assembly model, BBS5 assembles into the BBsome as a peripheral, non-framework subunit [10]. Novel BBs5 mutations are still being found eventually [23–25], but its actual function remains very limited understood. The two proteins appear to have been separated and independent, but the biological function they are both involved in show some potential coincidence. A systematic proteome-scale mappping has investigated ~14,000 binary protein-protein interactions by yeast-two-hybrid system. The huge dataset indicated an interaction between Klc3 and Bbs5 [23], which inspired us in further functional study on the relation of Klc3 and Bbs5. First of all, the protein-protein interaction was confirmed with Co-Immunopercipitation. Flag-tagged Klc3 and HA-tagged Bbs5 could pulled down each other (see Fig. 1), which means we already set a link between the Klc3 and BBsome. And we presumed that Klc3 might be functioning as a vehicle with the binding of the IFT Cargo complex, BBsome.

**FIGURE 1.** Co-immunoprecipitates of Klc3 with Bbs5, and Bbs4 with Bbs5. *A*, Immunoblot (IB) analysis of input cell lysates, anti-Flag immunoprecipitates (IP) derived from NIH3T3 cells transfected with pcDNA3-Flag-Kcl3 and pcDNA3-HA-Bbs5 constructs. *B*, Immunoblot (IB) analysis of input cell lysates, anti-Flag immunoprecipitates (IP) derived from NIH3T3 cells transfected with pcDNA3-Flag-Bbs4 and pcDNA3-HA-Bbs5 constructs.

Recent studies indicated that, in retina, Bbs5/Bbs5L (a newly found splice variant) localize at axonemes via binding to arrestin1 in a phosphorylation-dependent manner, and BBs5/Bbs5L are phosphorylated by protein kinase C (PKC) [15, 16]. The Bbs5 assembling properties in BBsome and the Bbs5-arrestin1 binding indicated that Bbs5 might be functioning as a docking compartment of BBsome, and the covalent modifications of phosphorylation might be a good entry point to investigate the BBS5 functions. For the time being, there is still no data about the PKC phosphorylation sites of Bbs5. As an important role in various biological processes, PKC also takes part in spermatogenesis [17–19]. Bbs5 might be one potential target of which PKC gets involved in since BBS5 is the subunit of BBsome, an indispensable complex during ciliogenesis and spermatogenesis [20, 21]. We analyzed Bbs5 as a potential PKC substrate in mice with the DISPHOS software [22], and two PKC phosphorylation sites, including Ser^19^ and Ser^246^, were predicted. But no data has been published about the exact phosphorylation sites. We performed the in vitro phosphorylation with purified Bbs5 protein as the substrate of PKC, and then the sample was analyzed with LC-MS/MS. Both Ser^19^ and Ser^246^ site were confirmed, consistent with DISPHOS prediction (see Fig. 2). The next question we need to figure out would be these sites function. We detected the endogenous Bbs5 expression in mouse brain and testis, with the customized Bbs5 phospho/nonphospho antibodies recognizing either site. Stronger phosphorylation of Bbs5-Ser^19^ was observed in both brain and testis. Interestingly, the phosphorylated Bbs5-Ser^246^ band was detected in brain but absent in testis (see Fig. 3. A). These results demonstrated that both Bbs5-Ser^19^ and Bbs5-Ser^246^ are phosphorylated in brain while only Bbs5-Ser^19^ in testis, suggesting that Bbs5 is phosphorylated in a tissue-specific pattern. And the dephosphorylation of Bbs5-Ser^246^ might play some role in Bbs5’s testis specific function. Also, we were interested in whether any PKC isoforms specifically phosphorylate Bbs5 in testis. The interactions were confirmed with Co-Ips, between 5 dominant PKC isoforms in testis, PKC α, β, γ, δ, ζ and Bbs5 while no significant difference was observed (see Fig. 3. B).

**FIGURE 2.**
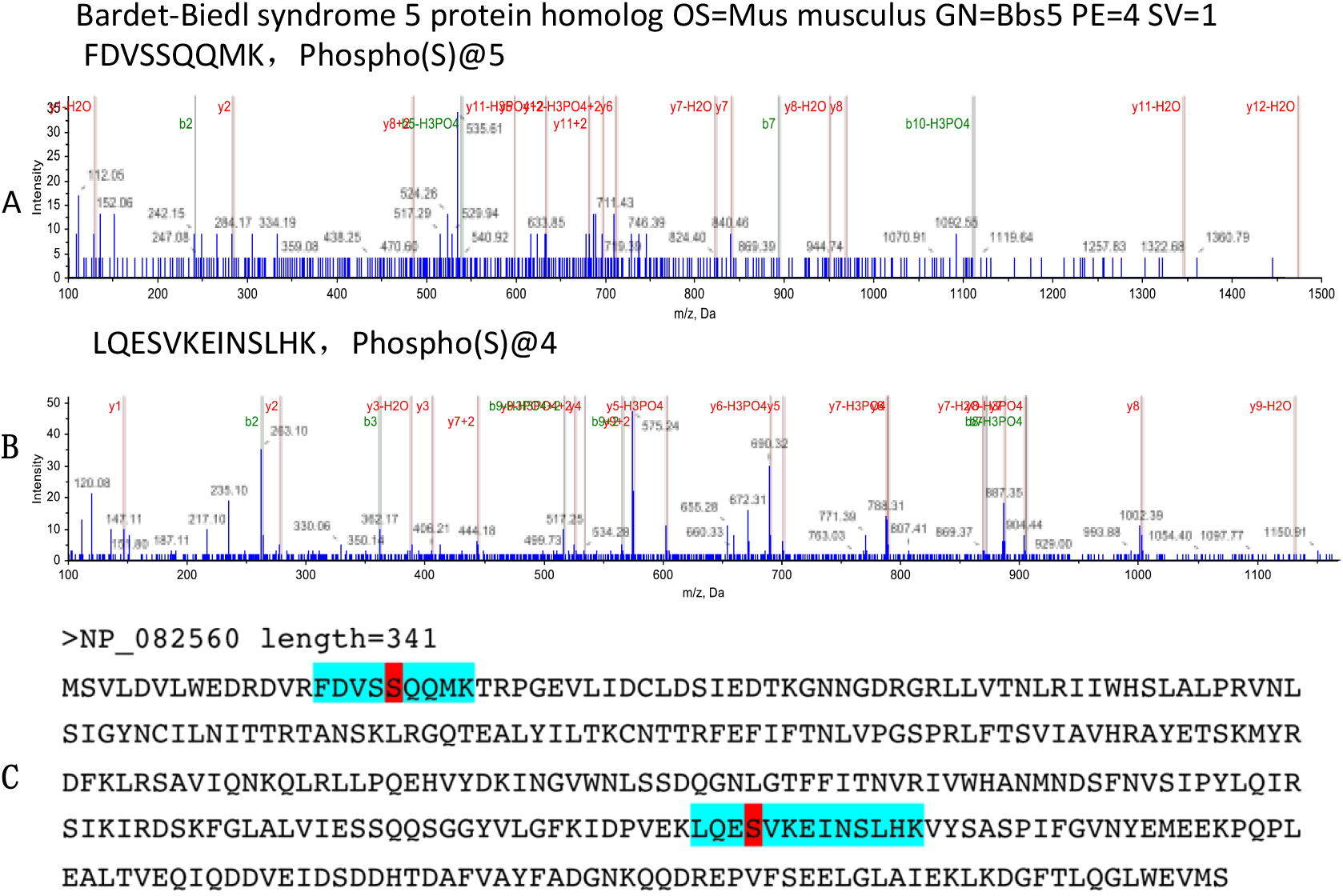
Bbs5 LC-MS/MS analysis identified two PKC phosphorylation sites, including Ser^19^ and Ser^246^. *A*, MS spectrum of the phosphorylated peptide, ^15^FDVSS^*^QQMK^23^, containing Ser^19^; S^*^ indicates that serine residue 19 is phosphorylated by PKC. *B*, MS spectrum of the phosphorylated peptide, ^242^LQES#VKEINSLHK^275^, containing Ser^246^, S# indicates neutral loss of H_3_PO_4_ from sequence ions. *C*, amino acid sequence of the Bbs5 protein. The LC-MS/MS analysis coverage rate of the given sample is >97% (the covered sequence is shown in *boldface* letters). The *color-highlighted* fragments indicate the identified phosphorylated peptide; _*p*_*S* represents phosphorylated serine, including Ser(P)^19^ and Ser(P)^246^.

**FIGURE 3.**
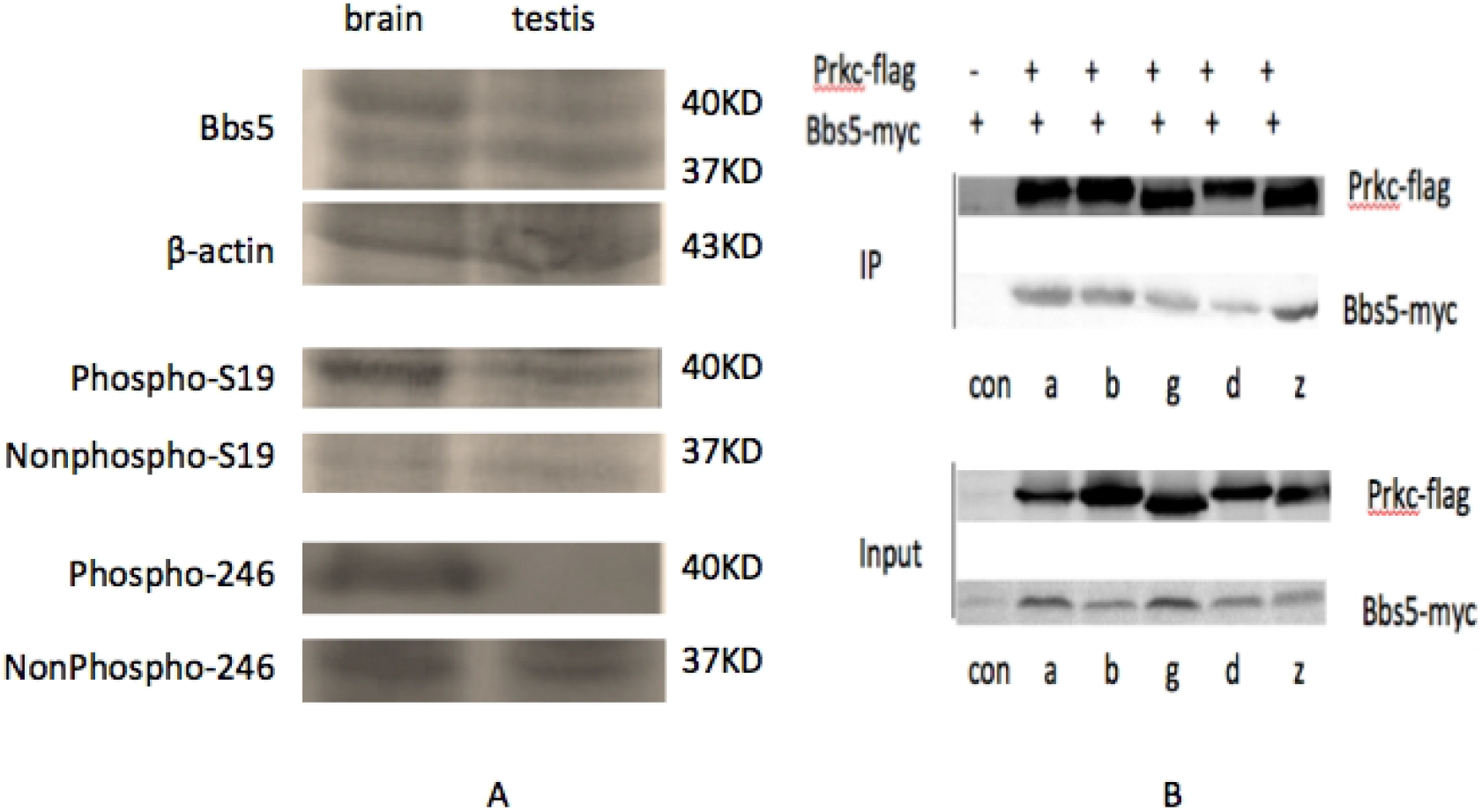
Endogenous Bbs5 protein expression and detection of the phosphorylation status of Bbs5-Ser^19^ and Bbs5-Ser^246^ in mouse brain and testis by Western blotting. *A*, phosphorylated Bbs5-Ser^19^ band was observed in both testis and S brain, whereas stronger phosphorylation of Bbs5-Ser^19^ was observed; a phosphorylated Bbs5-Ser^246^ band was detected in brain other than in testis. The molecular mass of proteins is indicated on the *right side* of the figure. ***B***, Immunoblot (IB) analysis of input cell lysates, anti-Flag immunoprecipitates (IP) derived from NIH3T3 cells transfected with Prkc-flag and Bbs5-myc constructs.

We further explored the role of Bbs5-Ser^19^ and Bbs5-Ser^246^ in regulating the Klc3-Bbs5 interaction. HA-Tagged De-phosphomimic and phosphomimic mutants constructs, Bbs5-S19A/S19D, Bbs5-S246A/S246D and Bbs5-S19A-S246A/S19D-S246D, were respectively co-overexpressed with Flag-tagged Klc3 at the present of PKC inhibitor H7 to down-regulate the endogenous phosphorylation. Surprisingly, Our findings showed functions of Bbs5-Ser^19^ and Bbs5-Ser^246^ in Klc3-Bbs5 interaction in diverse directions. Simulated phosphorylation/nonphosphorylation on Ser^19^ apparently have no effect on Bbs5-Klc3 interaction (see Fig. 4. A), suggesting that binding to Klc3 might be an intrinsic property of Bbs5 in mouse testis, regardless of Ser^19^ phosphorylation status; Interestingly, simulated phosphorylation on Ser^246^ completely block Bbs5-Klc3 interaction regardless of Ser^19^ (see Fig. 4. B & C), consistent with the tissue endogenous Bbs5 expression data (see Fig. 3. A).

**FIGURE 4.**
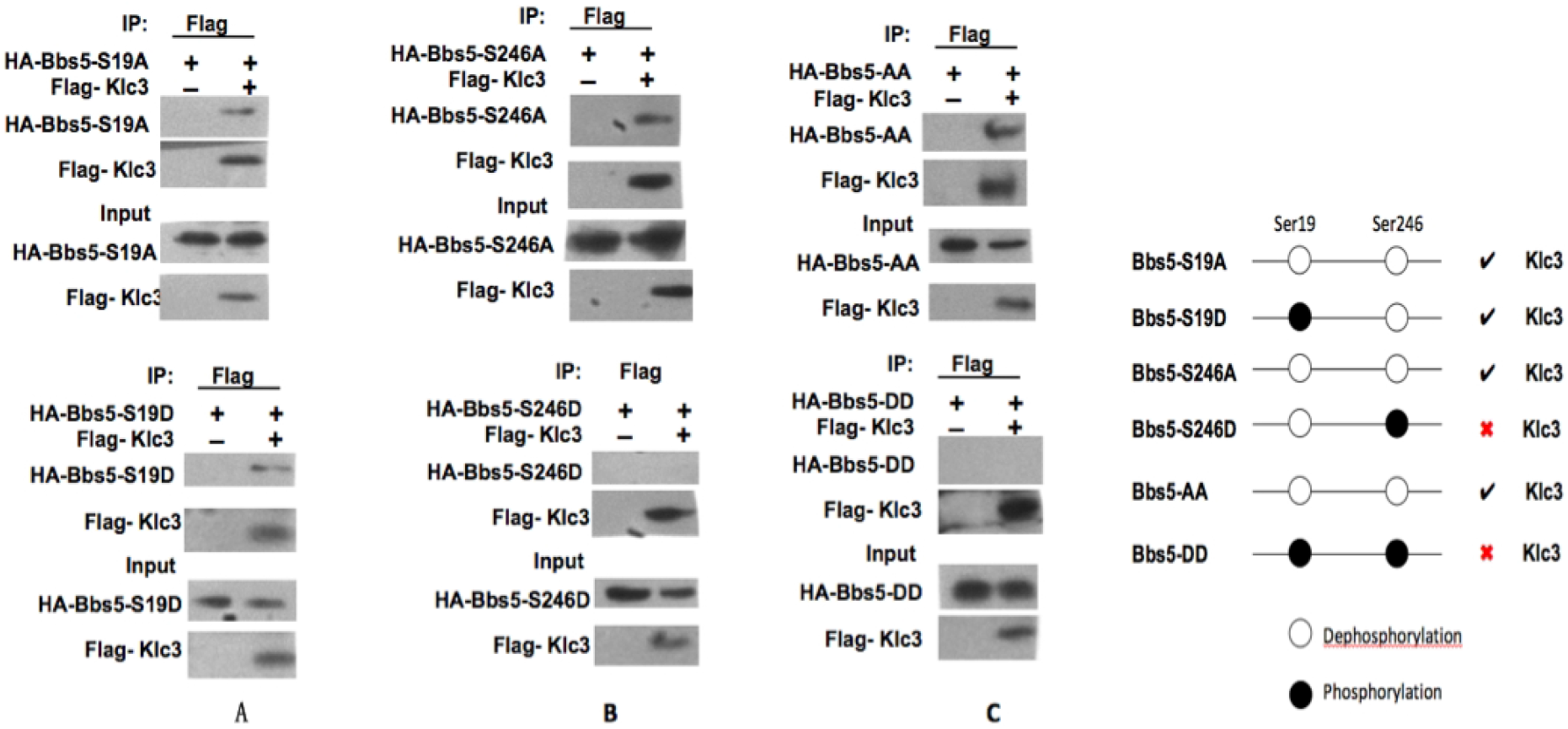
Co-IP of Klc3 with Bbs5 mutants, Bbs5-S19A, Bbs5-S246A, Bbs5-S19A/S246A, Bbs5-S19D, Bbs5-S246D, or Bbs5-S19D/S246D. *A*, Immunoblot (IB) analysis of input cell lysates, anti-Flag immunoprecipitates (IP) derived from NIH3T3 cells transfected with pcDNA3-Flag-Kcl3 and pcDNA3-HA-Bbs5 S19A/S19D constructs. *B*, Immunoblot (IB) analysis of input cell lysates, anti-Flag immunoprecipitates (IP) derived from NIH3T3 cells transfected with pcDNA3-Flag-Klc3 and pcDNA3-HA-Bbs5 S246A/S246D constructs. *C*, Immunoblot (IB) analysis of input cell lysates, anti-Flag immunoprecipitates (IP) derived from NIH3T3 cells transfected with pcDNA3-Flag-Klc3 and pcDNA3-HA-Bbs5 S19A+S246A/S19D+S246D constructs.

**FIGURE 5.**
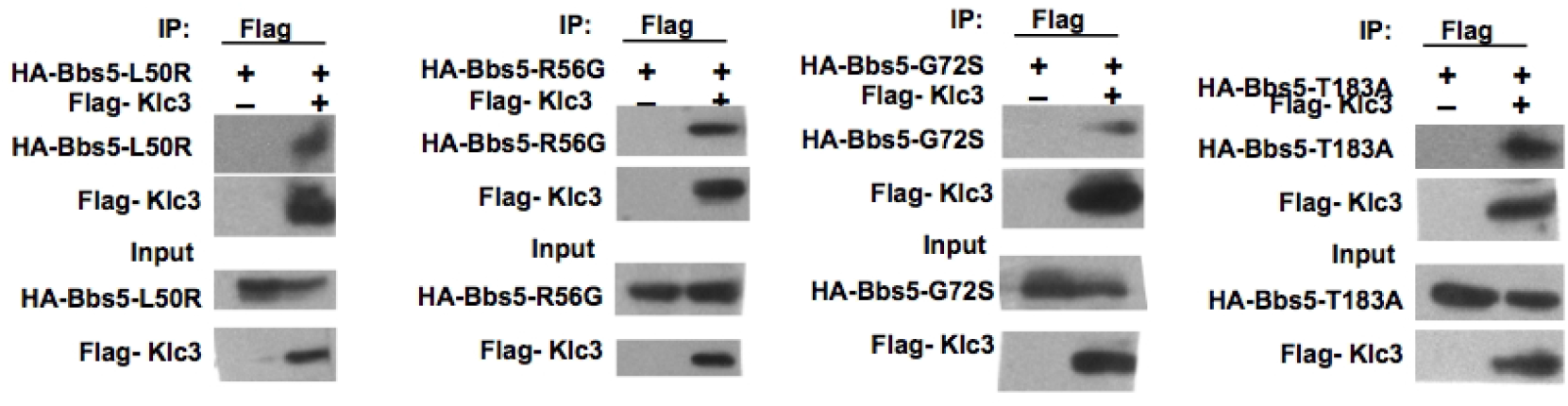
Co-IP of Klc3 with Bbs5 mutants, Bbs5-L50R, Bbs5-R56G, Bbs5-G72S and Bbs5-T183A. Immunoblot (IB) analysis of input cell lysates, anti-Flag immunoprecipitates (IP) derived from NIH3T3 cells transfected with pcDNA3-Flag-Klc3 and pcDNA3-HA-Bbs5 mutants constructs.

**FIGURE 5.**
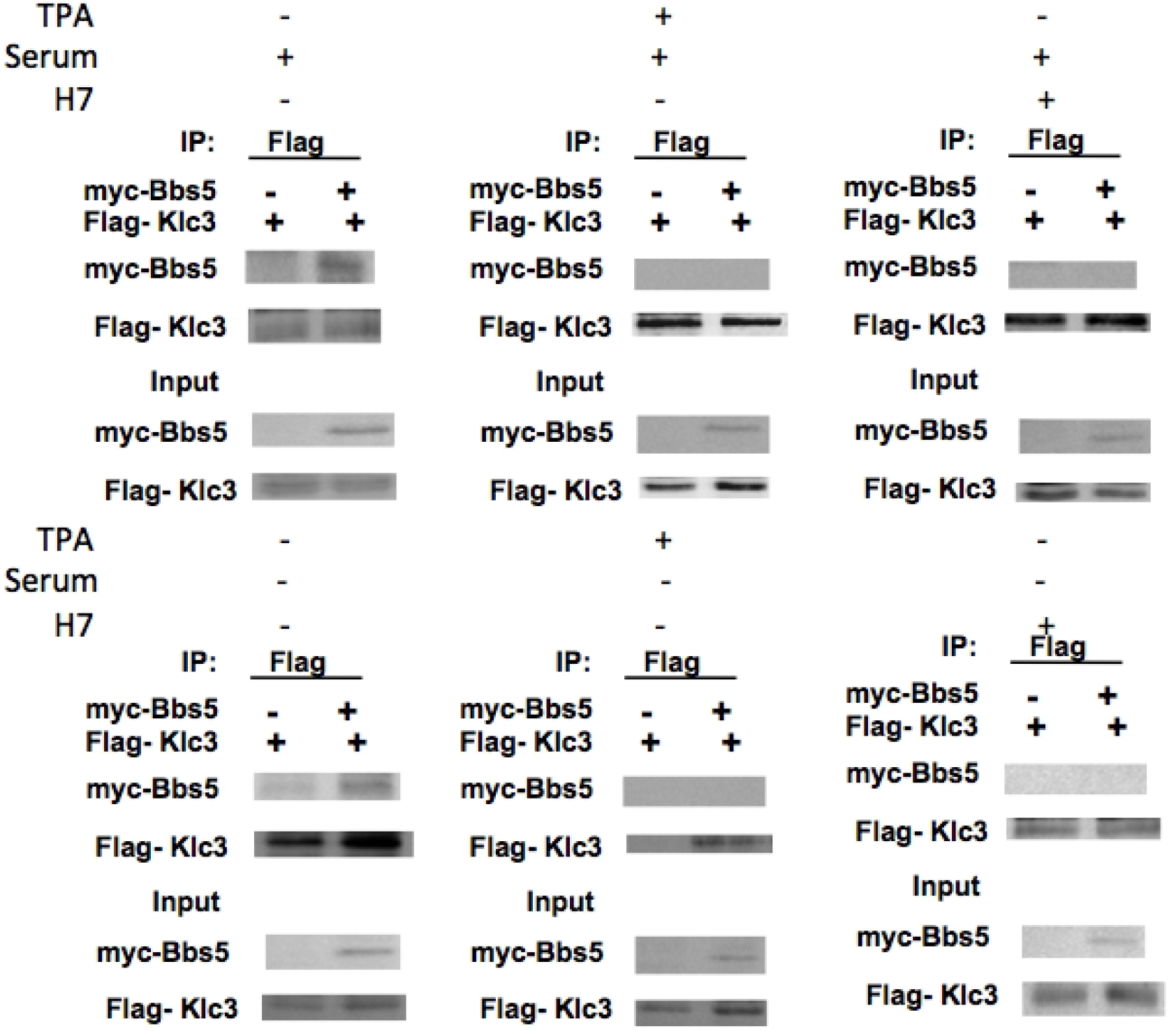
Klc3-Bbs5 Co-IP under different treatments. The upper row cells were cultured in medium with 10% serum, blank control, TPA and H7 were respectively added; The lower row cells were cultured in medium without serum, blank control, TPA and H7 were respectively added.

Additionally, We tried to observe whether several known BBS5 mutations could affect this interaction [23–25]. In this study, only the pathogenic missense mutants generating a full length Bbs5 protein were picked up for making the corresponding constructs. Co-IPs were performed and the data shows that none of mutations could 100% block the Klc3-Bbs5 interaction, while only G72S show a weakened pulled-down. More investigation needs to be done in future.

Our current results come to that the Klc3-Bbs5 binding in testis is more acting like the intrinsic assembly under the different phosphorylation status of Bbs5-Ser^19^ and Ser^246^. We also want to understand what if the PKCs were just diffusely stimulated or inhibited. The Klc3-Bbs5 binding was observed again under several different conditions. In the medium with/without serum, the PKC stimulator TPA or inhibitor H7 was applied to up-regulate or down-regulate the PKCs activity. Apparently, serum starvation itself does not block the binding. But surprisingly, both the stimulation and the inhibition of PKCs would almost 100% block the interaction, suggesting the diffuse stimulation and inhibition of PKCs might regulate Klc3-Bbs5 interaction in some alternative mechanism not fully elucidated. A hypothesis for right now would be the phosphorylation status of Bbs5-Ser^19^ and Ser^246^ were regulated by different PKC isoforms.

Collectively, the aforementioned findings could be accommodated in the following working model. In testis, the Bbs5 in fully assembled BBsome provides a docking site for Klc3. This docking site for Klc3 is only available in condition of Bbs5-Ser^246^ dephosphorylated. This testis-specific BBsome-Klc3 complex functions as a transportation cargo for the mitochondria to the designated subcellular localization. The temporary Bbsome-Klc3 complex releases Klc3 at the midpiece of sperm tail. Then Klc3 will continue executing other functions. And the BBsome keeps going along the flagellum.

In the future, it is of great importance to probe the exact mechanisms modulating the association of Klc3 with Bbs5, regulation of dephosphorylation of Bbs5-Ser^246^ and the other functions of diverse phosphorylation of Bbs5-Ser^19^ and Bbs5-Ser^246^ in various tissues.

## Experimental procedures

Kunming genealogy-specific pathogen-free mice (males at 8 weeks and ~30 g) were obtained from the Department of Laboratory Animals, China Medical University. All experiments were performed at China Medical University in accordance with the National Institutes of Health guidelines for the Care and Use of Laboratory Animals. Reagents, unless otherwise specified, were purchased from Sigma-Aldrich.

### Prokaryotic Expression and Purification of Recombinant Bbs5 Protein

A 1026-bp fragment encoding 341 amino acids was amplified by PCR with mouse testis cDNAs library as template, and PCR was performed with 35 cycles (primers: 5′-CGC<u>GGATCC</u>ATGTCTGTGCTGGACGTGTTG-3′ and 5′-CCG<u>CTCGAG</u>TCAACTCATCACTTCCCAAAGTC-3′). Each PCR cycle consisted of 30 s at 94 °C, 40 s at 64 °C, and 60 s at 72 °C. The PCR product was then subcloned into the pGEX-4T-2 vector. The recombinant plasmids, pGEX-4T-2-Bbs5-WT, was sequenced to verify the correct insertion.

The GST-Bbs5 was expressed for 3 h at 27 °C in *Escherichia coli*BL21(DE3) (Takara) transformed with pGEX-4T-2-Bbs5-WT in the presence of 0.1 mM isopropyl-β-D-thiogalactopyranoside (Takara). The proteins were purified with the glutathione-Sepharose 4B protein chromatography purification kit (GE Healthcare). Total protein was induced by isopropyl-β-D-thiogalactopyranoside, vector-expressed protein, and purified protein was identified with 10% SDS-PAGE and Western blotting.

### Phosphorylation of GST-Bbs5 Fusion Protein in Vitro and LC-MS/MS Analysis of PKC Phosphorylation Site

Purified GST-Bbs5 protein (400 μg) was incubated with 6 μl of PKC (New England Biolabs) and 10 μl of ATP (New England Biolabs) at 30 °C for 45 min. Following 10% SDS-PAGE separation, the band of phosphorylated Bbs5 protein was excised from the gel, and in-gel digestion was performed. To obtain maximal sequence coverage of protein, the digested products were collected at 1, 2, 4, and 12 h. The sample was combined and analyzed with LC-linear trap quadrupole-MS/MS (Thermo Fisher Scientific, Waltham, MA) with MS/MS (MS^2^) and MS/MS/MS (MS^3^) strategy. MS data were searched with SEQUEST, and the spectra of all identified phosphor-peptides were manually verified.

### Plasmid Construction and Site-directed Mutagenesis

A 1053-bp fragment encoding 5’-HA-tagged Bbs5 was amplified by PCR with pGEX-4T-2-Bbs5-WT as template, and PCR was performed with 35 cycles (primers: 5′-CAC<u>GGATCC</u>ATGTACCCATACGATGTTCCAGATTACGCT TCTGTG-3′ and 5′-CCG<u>CTCGAG</u>TCAACTCATCACTTCCCAAAGTC-3′). The PCR product was then subcloned into the pCDNA3 vector. The recombinant plasmids, pcDNA3-HA-Bbs5-WT, was sequenced to verify the correct insertion. The pcDNA3-HA-Bbs5-S19A construct was prepared by mutating Ser^19^ of Bbs5 to alanine with pcDNA-HA-Bbs5-WT as template and site-directed mutagenesis kit (Stratagene). The primers were FWD1 (5′-CTTCGACGTGTCCGCACAACAGATGAAAAC-3′) and REV2 (5′-GTTTTCATCTGTTGTGCGGACACGTCGAAG-3′). The pcDNA3-HA-Bbs5-S246A or pcDNA3-HA-Bbs5-S19A/S246A constructs were prepared by mutating Ser^246^ of Bbs5 to alanine with pcDNA3-HA-Bbs5-WT or pcDNA3-HA-Bbs5-S19A as templates. The primers were FWD3(5′-GTGGAAAAACTACAGGAAGCAGTTAAAGAGATCAACTCAC-3′) and REV4 (5′-GTGAGTTGATCTCTTTAACTGCTTCCTGTAGTTTTTCCAC-3′). Mutants mimicking phosphorylation, pcDNA3-HA-Bbs5-S19D, pcDNA3-HA-Bbs5-S246D and pcDNA-HA-Bbs5-S19D/S246D, were also constructed, in which residues 19 and/or 246 were replaced by aspartic acid. The primers were FWD5 (5′-CTTCGACGTGTCCGACCAACAGATGAAAAC-3′), REV6 (5′-GTTTTCATCTGTTGGTCGGACACGTCGAAG-3′), FWD7 (5′-GTGGAAAAACTACAGGAAGACGTTAAAGAGATCAACTCAC-3′) and REV8 (5′-GTGAGTTGATCTCTTTAACGTCTTCCTGTAGTTTTTCCAC-3′). With similar strategy, pcDNA3-myc-Bbs5-WT, pcDNA3-HA-Bbs5-L50R, pcDNA3-HA-Bbs5-R56G, pcDNA3-HA-Bbs5-G72S, pcDNA3-HA-Bbs5-T183A, pcDNA3-flag-Bbs4, pcDNA3-flag-Klc3 and pcDNA3-flag-PKC-α/β/γ/δ/ζ were also built up. All of the above recombinant plasmids were sequenced to verify the correct gene insertion and successful mutation.

### Western Blotting and Co-Immunoprecipitation

Mouse brain/testis and transfected NIH3T3 cells were lysed, subjected to SDS-PAGE (10%), transferred onto nitrocellulose membranes, and immunoblotted with antibodies to Bbs5 (1:200; Santa Cruz Biotechnology), phospho-Bbs5-pSer^19^ /nonphospho-Bbs5-Ser^19^, phospho-Bbs5-pSer^246^ /nonphospho-Bbs5-Ser^246^ (1:300; Bioyear Gene), GST/HA/Myc/Flag and β-actin (1:500; Beyotime Institute of Biotechnology). Proteins were visualized by the enhanced chemiluminescence (ECL) detection system (Pierce Biotechnology). The phospho-Bbs5-pSer^19^ /nonphospho-Bbs5-Ser^19^ and phospho-Bbs5-pSer^246^ /nonphospho-Bbs5-Ser^246^ antibodies were raised in New Zealand White Rabbits against the keyhole limpet hemocyanin-conjugated phosphopeptides, DRDVRFDVSpSQQMKTRPG / DPVEKLQEpSVKEINSLHK, or the nonphosphopeptides, DRDVRFDVSSQQMKTRPG / DPVEKLQESVKEINSLHK.

## Funding and additional information

* This work was supported by Liaoning Province Education Administration Doctoral Startup Fund Grant 201501011.

